# Homotypic SLAMF1:SLAMF1 interactions between innate T cells and neutrophils activate fungal killing by neutrophils

**DOI:** 10.64898/2026.02.19.706741

**Authors:** Lindsay S. Lau, Sarah Lichtenberger, Cleison Ledesma Taira, Bruce S. Klein, Marcel Wüthrich

**Affiliations:** Departments of Pediatrics; Internal Medicine; Medical Microbiology and Immunology University of Wisconsin & Infectious Diseases, Madison, WI, USA

## Abstract

Neutrophils and monocytes are the main fungal effector cells in restricting *Blastomyces dermatitidis* (*Bd)* and other fungi at the respiratory mucosa. However, understanding how phagocytes become activated and recruited to the site of infection is still incompletely understood. Innate lymphocytes and myeloid cells have been found to communicate and play an essential part in activating neutrophils and other effector cells to kill fungi. Here, we identified that Signaling Lymphocytic Activation Molecule 1 (SLAMF1) is a key host immune receptor involved in orchestrating a cellular and molecular signaling network that leads to the activation of phagocytes. By using mice to conditionally eliminate SLAMF1 receptor expression on innate CD4^+^ or TCRγδ^+^ T cells, we uncovered that these innate lymphocytes augment neutrophil killing of *Bd* in a SLAMF1 dependent manner. SLAMF1 expression on neutrophils enabled homotypic SLAMF1:SLAMF1 interactions with innate CD4^+^ T cells, which prompted release of soluble factors that activated neutrophils to kill fungi. Our work furnishes new mechanistic insight about the role of SLAMF1 in mobilizing innate immune cells to induce phagocyte-driven killing of inhaled fungi.

**Author Summary:** Emerging fungal diseases represent a significant and growing global public health threat fueled by increased anti-fungal resistance and rising number of immunocompromised individuals. Most fungal infections are respiratory and occur when inhaled fungal spores settle in the lungs and cause inflammation or tissue damage. The innate immune system is the first line of defense in the lungs but the mechanisms by which the host immune system becomes activated and mounts a protective response is still not completely understood. We identified a receptor on innate immune cells that facilitates communication between cells and recruits and activates killer cells that engulf and destroy the fungal pathogen. We uncovered that the receptor on the cell surface of innate immune cells mediates its function through cell contact and induction of soluble factors. Our work offers new mechanistic insight about how the innate immune system becomes activated by the presence of fungi and orchestrates an effective host response. We envision that soluble receptor could be harnessed for future anti-fungal therapy.

## INTRODUCTION

Antimicrobial defense against fungal pathogens requires the concerted activation of a signaling network of lung epithelial cells, innate lymphocytes, and myeloid cells. Crosstalk between myeloid and lymphoid cells help control inhaled fungi [1,2]. Upon fungal infection, innate lymphocytes including CD4^+^TCRβ^+^, TCRγδ^+^, MAIT, NK, NKT and innate lymphoid cells become activated and produce soluble factors that activate phagocytes such as neutrophils, monocytes and macrophages to kill fungi [3,4,5]. These studies have clarified the cells that initially respond to inhaled fungi but have not defined the cell surface receptors that mediate activation and communication between these cells.

Innate immune responses to microbial pathogens are orchestrated by interaction between cell surface molecules expressed on leukocytes. Signaling Lymphocytic Activation Molecule Family (SLAMF) receptors play a crucial role in regulating and interconnecting both innate and adaptive immune cells [6]. SLAMF receptors are widely expressed on hematopoietic lineage cells [7] and have three modes of action: (i) homophilic (receptor-to-receptor) contact-dependent interaction between cells that modulate immune responses (*e.g.* costimulatory interactions between DC and T cells); (ii) direct microbial sensing as part of the anti-microbial response; and (iii) serving as an entry receptor for pathogens (*e.g.* for measles virus) [8,9,10]. The role of SLAMF1 in antifungal resistance is important but understudied. From our published work [11], we have learned that SLAMF1 is dispensable in vaccine induced adaptive immunity, but essential for innate defenses against fungal infection. Since SLAMF1 is of paramount importance in antifungal host defense [11], understanding its role in the activation and signaling of innate immunity at lung mucosa represents a gap in knowledge. Our premise is that SLAMF1 on hematopoietic cells plays a direct role in activating phagocytes to eliminate fungal and other microbial pathogens. We hypothesize that SLAMF1 mediates neutrophil activation through homophilic (e.g. SLAMF1-to-SLAMF1) signaling interactions between SLAMF1 receptor expressing innate lymphocytes and myeloid cells. To test our hypothesis, we generated conditional SLAMF1 knockout mice that lack the receptor in candidate cells for activating neutrophils during fungal infection to evaluate the role of SLAMF1 in restraining fungal growth *in vivo* and *in vitro*.

Here, we report that 1) SLAMF1 is required for the innate immune response to pulmonary infection with *Bd*, *in vivo* killing of yeast by neutrophils and monocytes, and ROS and NO production by the cells, 2) innate CD4^+^ T cells, TCRγδ^+^ T cells, MAIT cells, and monocytes express SLAMF1 following pulmonary fungal infection, 3) SLAMF1 on innate CD4^+^ T cells is indispensable for restraining pulmonary fungal infection, whereas SLAMF1 is dispensable on TCRγδ^+^ T cells and monocytes, 4) SLAMF1 expressing innate CD4^+^ T cells augment neutrophil killing through cognate, homotypic SLAMF1:SLAMF1 interactions with neutrophils and non-cognate, soluble factors. Our studies clarify how SLAMF1 facilitates crosstalk between hematopoietic cells and enables neutrophils to restrain fungal infection.

## RESULTS

### SLAMF1 is required for innate resistance to Bd and in vivo killing by neutrophils and monocytes

We recently reported that vaccinated SLAMF1 knockout mice fully develop antigen-specific CD4^+^ T cells that migrate to the lung upon challenge as with vaccinated wild type mice. Although antigen-specific CD4^+^ T cells protected vaccinated wild type mice, they failed to protect vaccinated SLAMF1 knockout mice [12]. These results implied that innate resistance to *Bd* infection might depend on SLAMF1. Indeed, unvaccinated SLAMF1 knockout mice died earlier (**Fig. 1A**) and had 3,500-fold more lung CFU than wild-type mice at a time when knockout mice were moribund (**Fig. 1B**). These findings pointed to a deficit in early, innate resistance to fungal infection in SLAMF1 knockout mice.

**Figure 1:**
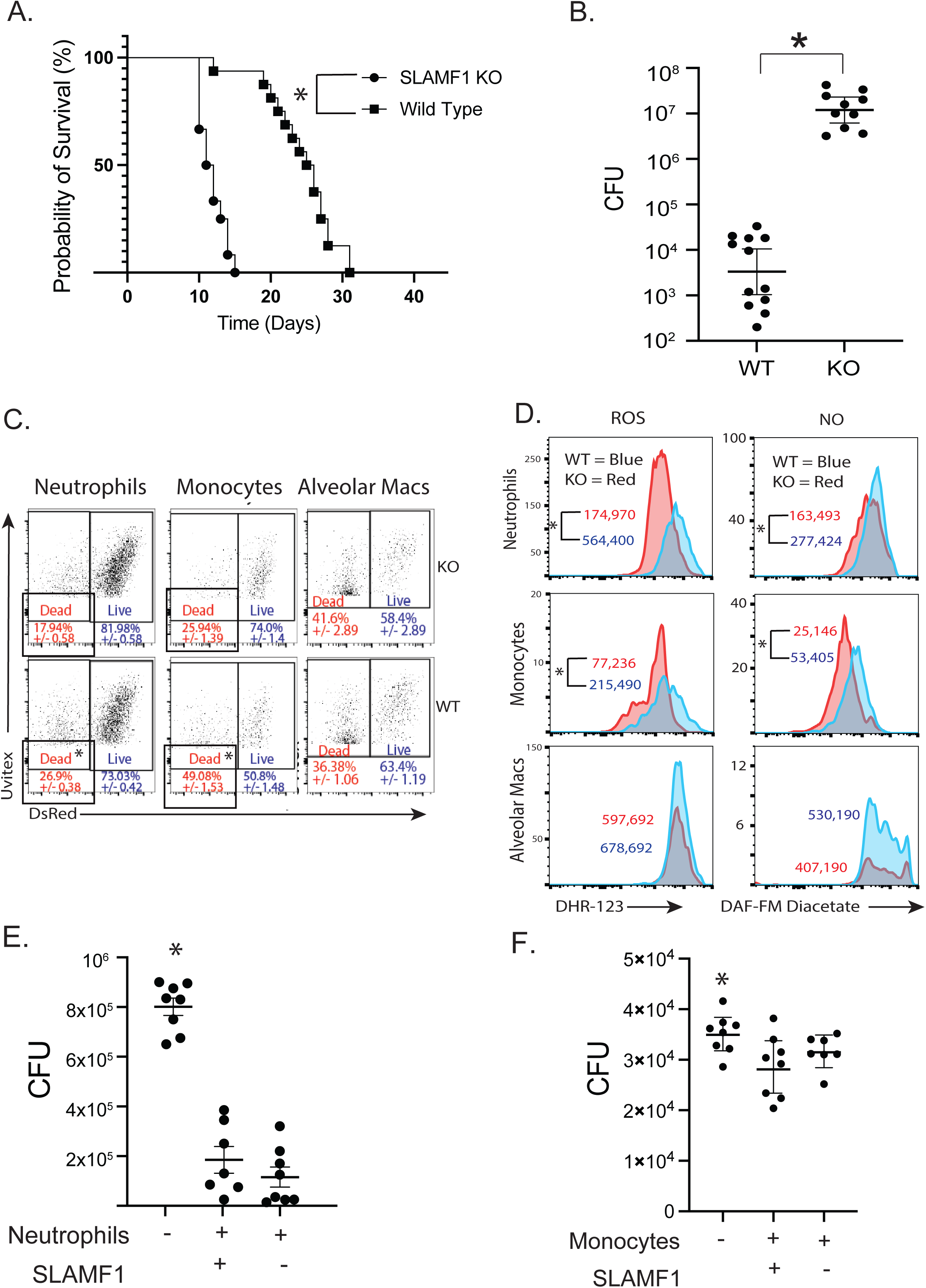
SLAMF1 is required for innate resistance to *Bd*, *in vivo* killing and ROS/NO production by neutrophils and monocytes. Survival **(A)** and lung CFU **(B)** 4 days post infection of SLAMF1 knockout (KO) and wild type (WT) mice infected with *Bd*. *p<0.05, Kaplan Meier test for survival. CFU from at least 10 mice/group are expressed as Log_10_ plotted with geometric mean ± geometric SD *p<0.05, two tailed Mann-Whitney T test. *In vivo* killing assay: Dot plots show yeast-associated cells (alveolar macrophages, neutrophils and monocytes) and the percentages of cells containing dead (DsRed-, Uvitex+) vs. live (DsRed+, Uvitex+) yeast. Dot-plots are concatenated data for 5 mice/group *p<0.05 vs KO, two tailed Mann-Whitney T test **(C)**. Lung cells stained *ex vivo* with ROS or NO indicators DHR-123 or DAF-FM diacetate. Geometric MFI of staining is shown for yeast-associated neutrophils, monocytes, and alveolar macrophages *p<0.05 two tailed Mann-Whitney T test **(D**). *In vitro* killing assay: neutrophils and monocytes were isolated from bone marrow of wildtype and SLAMF1 KO mice and cocultured with *Bd* yeast overnight at 37C. 8 replicates plated for CFU. *p<0.05 vs. all other groups, two tailed Mann-Whitney T test **(E+F)**.

Phagocytes such as neutrophils, monocytes, and macrophages take up and kill fungi [13]. To see if SLAMF1 impacts elimination of *Bd* by phagocytes *in vivo*, we used a fluorescent DsRed viability reporter strain that we engineered [14,15]. The strain “reports” *in vivo* killing as measured by loss of red fluorescence. We assessed killing by neutrophils, monocytes and alveolar macrophages at 16 hours post-infection when the burden of lung CFU was comparable between wild type and SLAMF1 knockout mice.

We found that the percentage of dead yeast in neutrophils and monocytes, but not alveolar macrophages is reduced in SLAMF1 knockout mice compared to wild-type mice (**Fig. 1C + SFig. 1A**). Thus, killing of *Bd* by neutrophils and monocytes is impaired in the absence of SLAMF1 [13]. While the percentage of killed yeast associated with neutrophils (26.9%) is lower than with monocytes (49.08%) the overall number of yeast-associated neutrophils is 4.3 times higher than yeast associated monocytes (**SFig. 1B**) indicating that innate resistance to fungal infection is mostly determined by killing of yeast by neutrophils. Thus, we focused our study on the activation of neutrophils to understand how SLAMF1 promotes innate host resistance.

Production of reactive oxygen species (ROS) and/or nitric oxide (NO) is essential and serves as a valid surrogate of innate resistance to many fungi [14]. We investigated whether reduced *in vivo* killing by neutrophils in SLAMF1 knockout mice is associated with reduced production of these products. We found that staining of ROS and NO was reduced in yeast-associated neutrophils and monocytes in SLAMF1 knockout vs wild type mice (**Fig. 1D**). Consistent with intact *in vivo* killing by alveolar macrophages, SLAMF1 did not affect the production of ROS and NO in these cells.

To assess intrinsic killing by SLAMF1-deficient neutrophils and monocytes, we cocultured neutrophils or monocytes with *Bd in vitro* and measured killing. In contrast to the *in vivo* defect, *in vitro* killing of *Bd* was similar by SLAMF1 deficient and SLAMF1 sufficient neutrophils (**Fig. 1E**) and monocytes. **(Fig. 1F)**. These results indicate that neutrophils and monocytes from SLAMF1-deficient mice did not exhibit an intrinsic defect in killing *Bd* yeast and that, *in vivo,* SLAMF1 is required on other cells for antifungal function of neutrophils and monocytes. Thus, we sought to identify the innate leukocytes that express SLAMF1 to promote antifungal actions of neutrophils and monocytes.

### Innate lymphocytes and monocytes express SLAMF1

To identify potential immune cells that could mediate SLAMF1-dependent activation of phagocytes, we stained lung cells with anti-SLAMF1 (CD150) monoclonal antibody at 16 hours post-infection when the lung burden of fungal infection is still comparable between SLAMF1 knockout and wild-type mice. We used a panel of multiplexing antibodies to identify 17 pulmonary leukocyte subsets with 12 color flow cytometry [16]. Among the 17 populations of myeloid and lymphoid cells, we found three innate lymphocyte subsets (CD4^+^TCRβ^+^, TCRγδ^+^, MAIT cells) and one myeloid cell population (Ly6C^hi^ CCR2^+^ monocytes) that expressed SLAMF1 in wild type mice but not in SLAMF1 knockout mice (**Fig. 2**). Below we systematically investigated whether CD4^+^TCRβ^+^, TCRγδ^+^, and Ly6C^hi^ CCR2^+^ monocytes activate neutrophils in a SLAMF1-dependent manner. MAIT cells were too few in numbers in the lung after fungal infection and there aren’t any MAIT cell specific cre mice available to study the role of SLAMF1 on MAIT cells.

**Figure 2:**
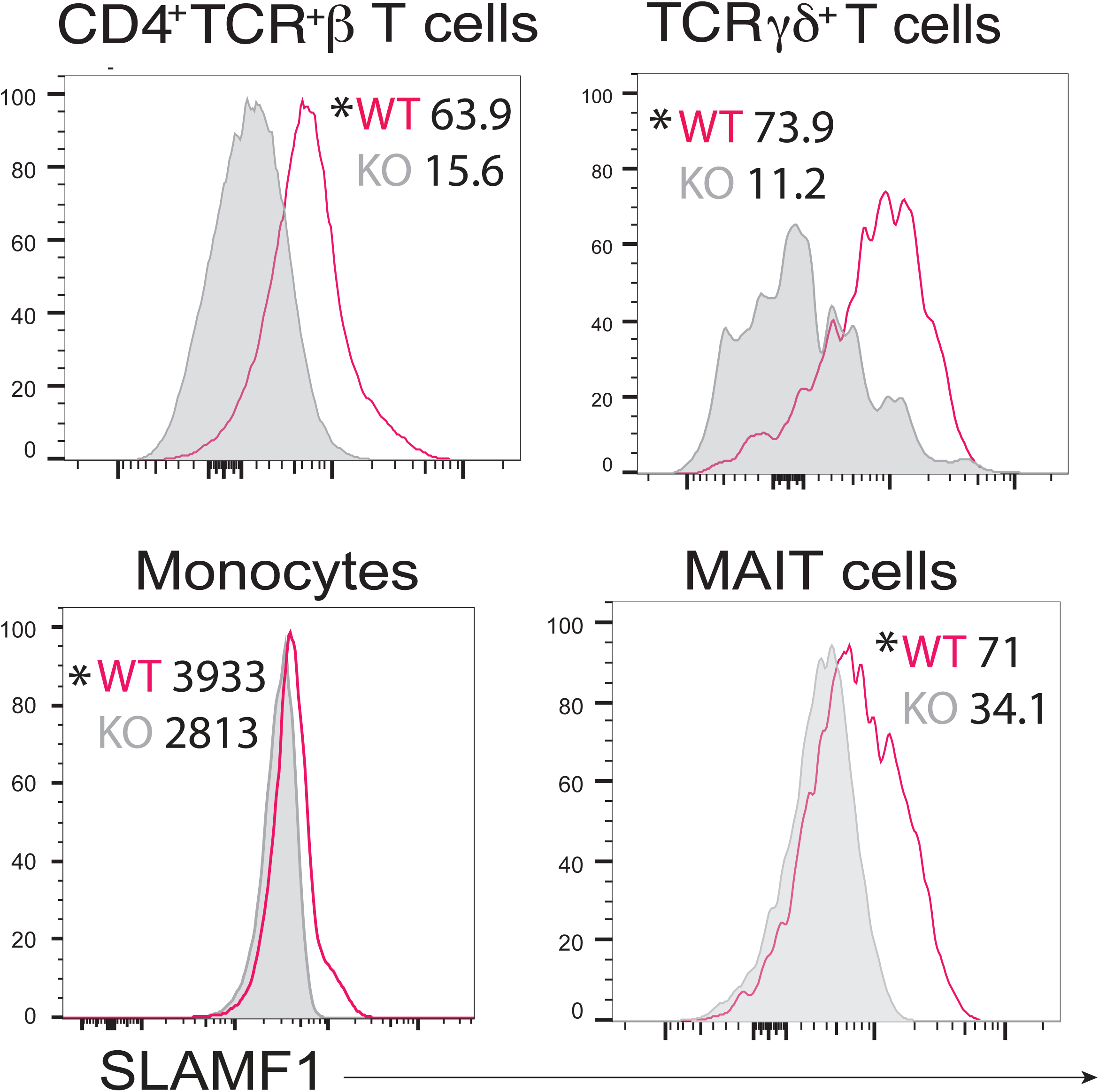
Innate lymphocytes and monocytes express SLAMF1. *Ex vivo* staining of lung leukocytes for SLAMF1. WT and SLAMF1 KO mice were challenged with *Bd* and lungs harvested at 16 hr post-infection. Lung cells were stained *ex vivo* with anti-SLAMF1 (CD150) mAb. SLAMF1 staining of cells from WT and SLAMF1 KO mice (shaded. The plots are concatenates from 5 mice/group. P values were calculated based on individual sample MFIs between WT and SLAMF1KO mice. p<0.05 vs. KO for all cell types, two tailed Mann-Whitney T test

### Innate CD4^+^ T cells, TCRγδ^+^ T cells and monocytes augment neutrophil killing of Bd in vitro

To determine whether SLAMF1 expressing lymphocytes or monocytes activate neutrophils to increase fungal killing, we established an *in vitro* coculture assay with yeast, neutrophils, and the candidate cell types (**Fig. 3A**). We cocultured *Bd* yeast with naïve neutrophils from the bone marrow and SLAMF1 expressing cells harvested from the lungs 16 hours after challenge. The purity of isolated cells was >90% for CD4^+^ T cells and monocytes, and 56% for TCRγδ^+^ T cells **(SFig. 2).** After coculture, neutrophils alone reduced yeast numbers **(Fig 3B-D).** The addition of CD4^+^ T cells, TCR-γδ^+^ T cells or monocytes further reduced CFU **(Fig 3B-D)**. These results suggested that CD4^+^ T cells, TCRγδ^+^ T cells and monocytes augment fungal killing by neutrophils.

**Figure 3:**
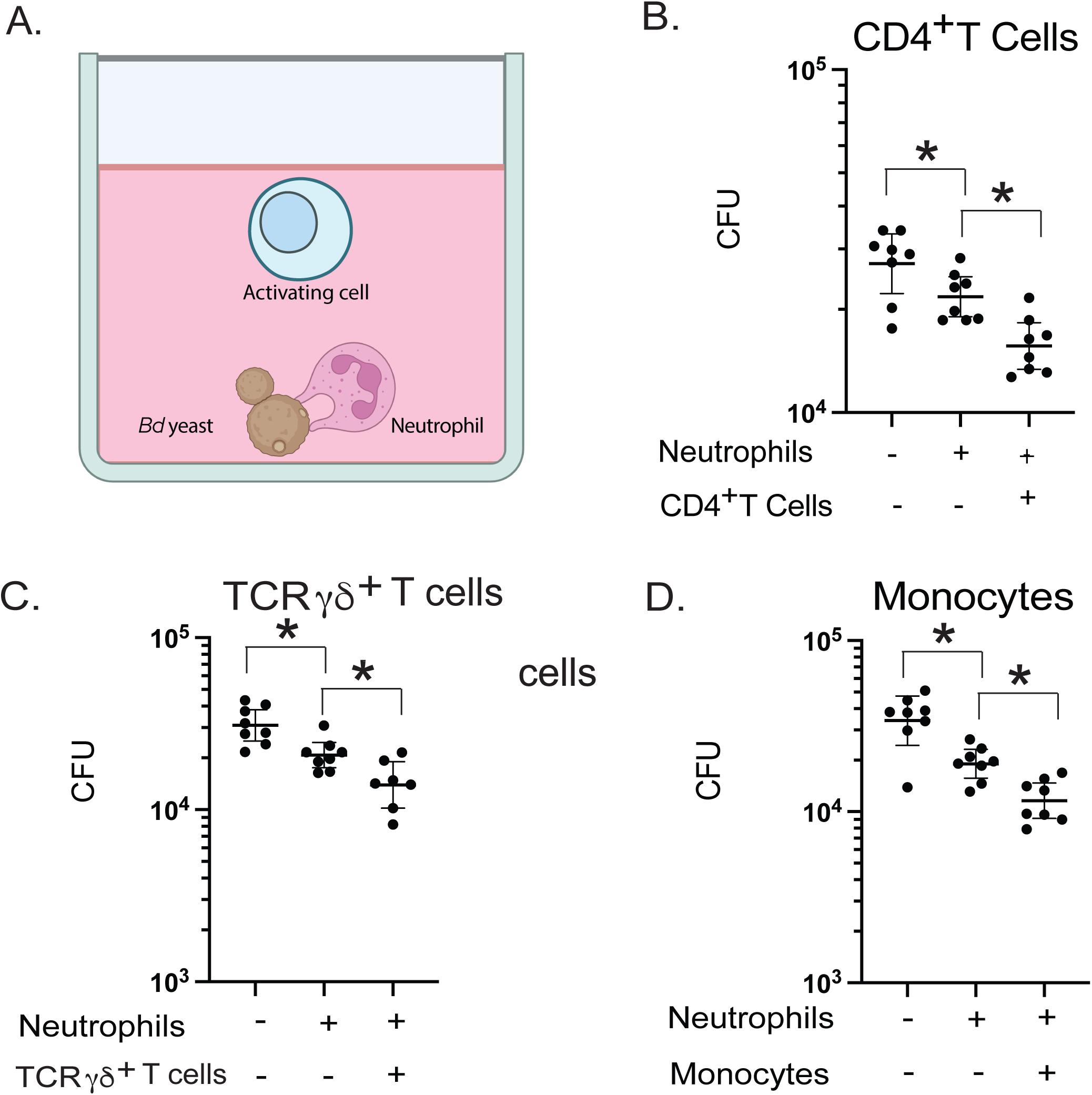
CD4^+^ T cells, TCRγδ^+^ T cells, and monocytes augment neutrophil killing of Bd. **(A)** *In vitro* killing assay with candidate activating cells, neutrophils and *Bd* yeast. Neutrophils were isolated from bone marrow of WT mice. Candidate cells were isolated from lungs of WT mice 16 hr after challenge. Neutrophils plus CD4^+^ T cells **(B)**, TCRγδ^+^ cells **(C)** and monocytes **(D)** were cocultured with *Bd* yeast overnight. 8 replicates plated for CFU. *p<0.05 two tailed Mann-Whitney T test. Bars are geometric mean with 95% CI. Graphs representative of 5 (B), 2 (C), and 2 (D) experiments, respectively.

### Generation and validation of conditional KO mice that lack SLAMF1on innate lymphocytes or monocytes

To determine whether augmented killing by added SLAMF1 expressing cells is truly dependent on SLAMF1 and whether SLAMF1 expressed by candidate cells is dispensable for host resistance to fungal infection *in vivo*, we generated conditional knockout mice in which SLAMF1 is ablated in the target cell population by Tamoxifen treatment. We validated the commercial SLAMF1 floxed mice by generating Ella cre x SLAMF1^fl/fl^ mice that lacked SLAMF1 expression in embryonic cells and phenocopied SLAMF1 cell surface staining and resistance to *Bd* infection of SLAMF1global knockout mice (**SFig. 3**).

To assess the role of SLAMF1 on individual target cells we crossed SLAMF1*^fl/fl^* mice with target cell specific Cre^+^ mice as outlined above (**SFig. 3**). Upon tamoxifen treatment we verified the loss of SLAMF1 expression on the target cells by FACS. 16 hours post-infection, CD4^+^ T cells from the lungs of CD4-Cre^+^ x SLAMF1*^fl/fl^* mice exhibited reduced SLAMF1 staining compared to CD4-Cre^-^ x SLAMF1*^fl/fl^* mice and wild type mice, similar to full body SLAMF1 knockout mice (**SFig. 4A**). We also observed a loss of SLAMF1 expression by TCRγδ^+^ T cells from TCRγδ-Cre^+^ x SLAMF1*^fl/fl^* mice (**SFig. 4B**) and monocytes from CCR2-Cre^+^ x SLAMF1*^fl/fl^* mice (**SFig. 4C**). SLAMF1 expression on monocytes was modest, so we used GFP expression as a surrogate marker to verify Cre activation and elimination of SLAMF1. Tamoxifen treatment of CCR2-Cre^+^ x SLAMF1*^fl/fl^* mice induced GFP expression on these cells (**SFig. 4D**); treatment of CCR2-Cre^-^ x SLAMF1*^fl/fl^* control mice did not.

### CD4^+^ T cells and TCRγδ^+^ T cells augment neutrophil killing in a SLAMF1-dependent manner

Since multiple SLAMF1 expressing cells (CD4^+^ T cells, TCRγδ^+^ T cells and monocytes) augmented fungal killing by neutrophils (**Fig. 3**), we sought to investigate whether each SLAMF1 expressing cell augments neutrophil killing in a SLAMF1-dependent manner. To address this question, we cocultured yeast and neutrophils with candidate activating cells from Cre^+^ and Cre^-^ conditional knockout mice (**Fig. 4A**). The addition of CD4^+^ T cells from CD4-Cre^-^ x SLAMF1*^fl/fl^* mice (express SLAMF1 by CD4^+^ T cells) but not CD4-Cre^+^ x SLAMF1*^fl/fl^* mice (lack SLAMF1 expression by CD4^+^ T cells) reduced CFU compared to neutrophils alone with yeast (**Fig. 4B**). Similarly, the addition of TCRγδ^+^ T cells from TCRγδ-Cre^-^ x SLAMF1*^fl/fl^* mice but not TCRγδ-Cre^+^ x SLAMF1*^fl/fl^* mice augmented neutrophil killing compared to neutrophils and yeast alone **(Fig. 4C).** These results indicate that both CD4^+^ T cells and TCRγδ^+^ T cells augmented killing by neutrophils in a SLAMF1-dependent manner.

**Figure 4.**
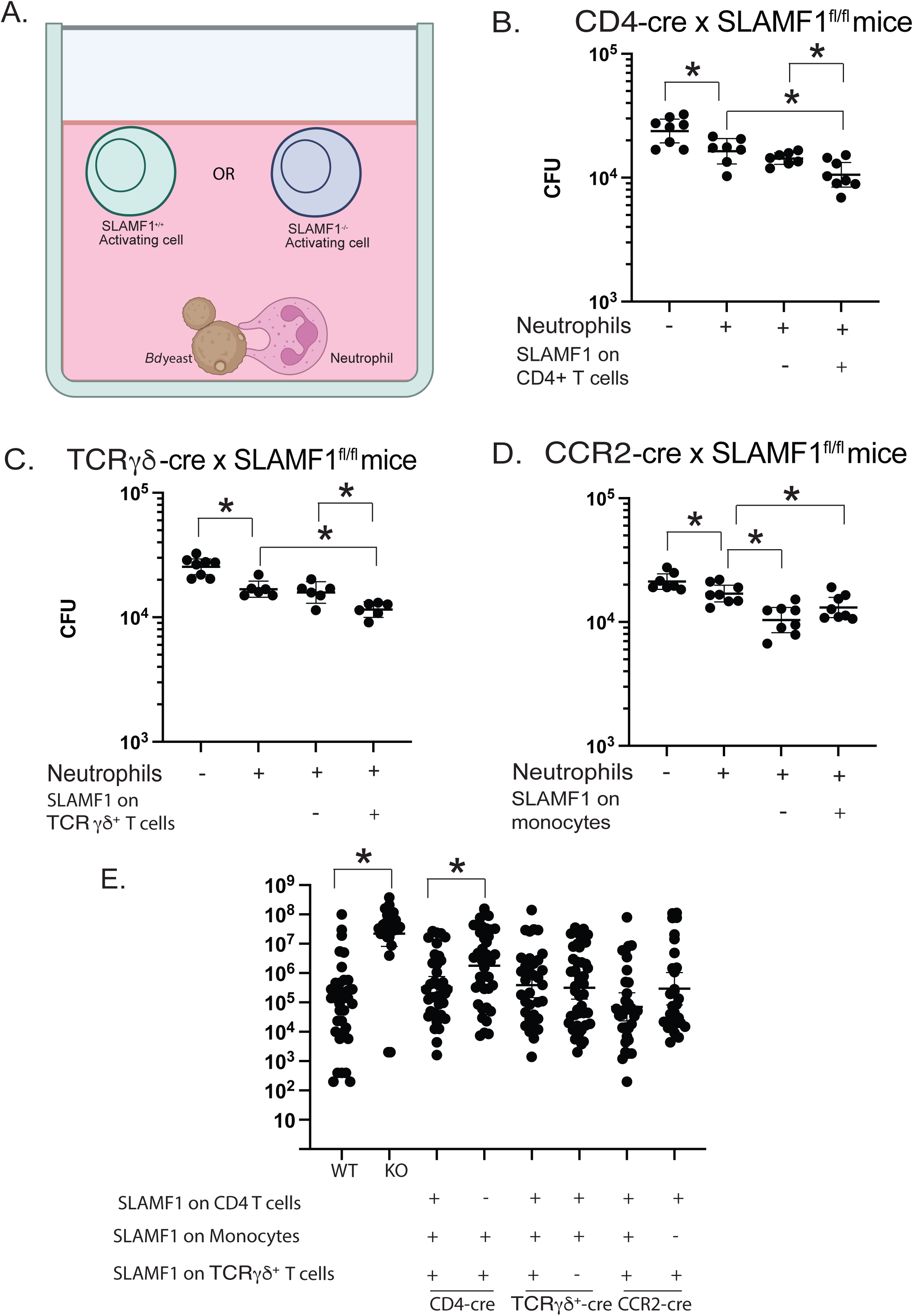
CD4^+^ T cells and TCRγδ^+^ T cells augment neutrophil killing in a SLAMF1-dependent manner and *in vivo* killing of *Bd* in conditional SLAMF1knockout mice. *In vitro* killing assay with candidate activating cells from conditional SLAMF1 knockout mice **(A)**. Neutrophils isolated from bone marrow of wild type mice were cocultured with *Bd* yeast and candidate activating cells from lungs of conditional CD4-Cre x SLAMF1^fl/fl^ **(B)**, TCRγδ-Cre x SLAMF ^fl/fl^ mice **(C)**, and CCR2-cre SLAMF1^fl/fl^ mice **(D)** harvested 16 hr post-infection. 8 replicates plated for CFU. Data representative of 3 assays. *p<0.05, two tailed Mann-Whitney T test. Wild type (WT), SLAMF1 knockout (KO), conditional CD4-cre x SLAMF1^fl/fl^ mice, CCR2-cre x SLAMF1^fl/fl^ mice, and TCRγδ-cre x SLAMF1^fl/fl^ mice were infected with 2×10^4^ *Bd* yeast and lung CFU plated at 14 days post-infection. Cre^+^ mice lack SLAMF1 on corresponding conditional knockout mice, Cre^-^ mice served as wildtype littermate controls for the corresponding Cre^+^ knockout mice **(E)**. *p<0.05 two tailed Mann-Whitney T test. Bars represent geometric mean with 95% CI.

In contrast, we did not find a difference in CFU reduction by adding monocytes from CCR2-Cre^-^ x SLAMF1*^fl/fl^* mice (express SLAMF1 on monocytes) or CCR2-Cre^+^ x SLAMF1*^fl/fl^* mice (lack SLAMF1 on monocytes) (**Fig. 4D**). These results indicate that reduced CFUs by the addition of monocytes is likely due to the ability of monocytes to also kill yeast rather than augmenting function/killing by neutrophils in a SLAMF1 dependent manner.

### SLAMF1 on CD4^+^ T cells is indispensable for anti-fungal resistance in vivo

To determine dispensability of SLAMF1 on target cells to host resistance to primary infection *in vivo*, we challenged the conditional knockout mice and analyzed lung CFU at the time these mice became moribund. Similar to full body SLAMF1 knockout mice, CD4-Cre^+^ x SLAMF1*^fl/fl^* mice that lacked SLAMF1 on CD4^+^ T cells had increased lung CFU compared to CD4-Cre^-^ x SLAMF1*^fl/fl^* control mice and wild type mice (**Fig. 4E**).

These results indicate that SLAMF1 expression on CD4^+^ T cells is indispensable for host resistance during primary infection. In contrast, lung CFU between TCRγδ-Cre^+^ x SLAMF1*^fl/fl^* mice and TCRγδ-Cre^-^ x SLAMF1*^fl/fl^* mice and CCR2-Cre^+^ x SLAMF1*^fl/fl^* mice and CCR2-Cre^-^ x SLAMF1*^fl/fl^* mice showed no significant difference (**Fig. 4E**). These results suggest that SLAM1 on TCRγδ-T cells and monocytes is dispensable *in vivo*, but it does not exclude a contribution for host resistance to primary infection in wild type mice.

### Augmented neutrophil killing requires SLAMF1 expression on neutrophils

The addition of CD4^+^ T cells and TCRγδ^+^ T cells augmented neutrophil killing in a SLAMF1 dependent manner *in vitro*. We sought to dissect the mechanism by which SLAMF1 increased neutrophil killing of *Bd* yeast. Our results above implied a homotypic interaction. Thus, if neutrophil activation by innate lymphocytes is mediated by SLAMF1:SLAMF1 homotypic interactions, we would expect that SLAM1 expression is required on both innate lymphocytes and neutrophils. To test this hypothesis, we established an *in vitro* coculture assay with CD4^+^ T cells and neutrophils from wild type or SLAMF1 deficient mice. The addition of neutrophils alone from wild type or SLAMF1-deficient mice reduced the number of yeast compared to coculture of yeast alone, confirming that SLAMF1-deficient neutrophils have no intrinsic defect in killing yeast.

The addition of CD4^+^ T cells from wild type mice reduced CFU in the wells with neutrophils from wild type but not SLAMF1-deficient mice (**Fig. 5A**). These results indicate that homotypic SLAMF1 interaction between neutrophils and CD4^+^ T cells is required to augment killing by neutrophils.

**Figure 5.**
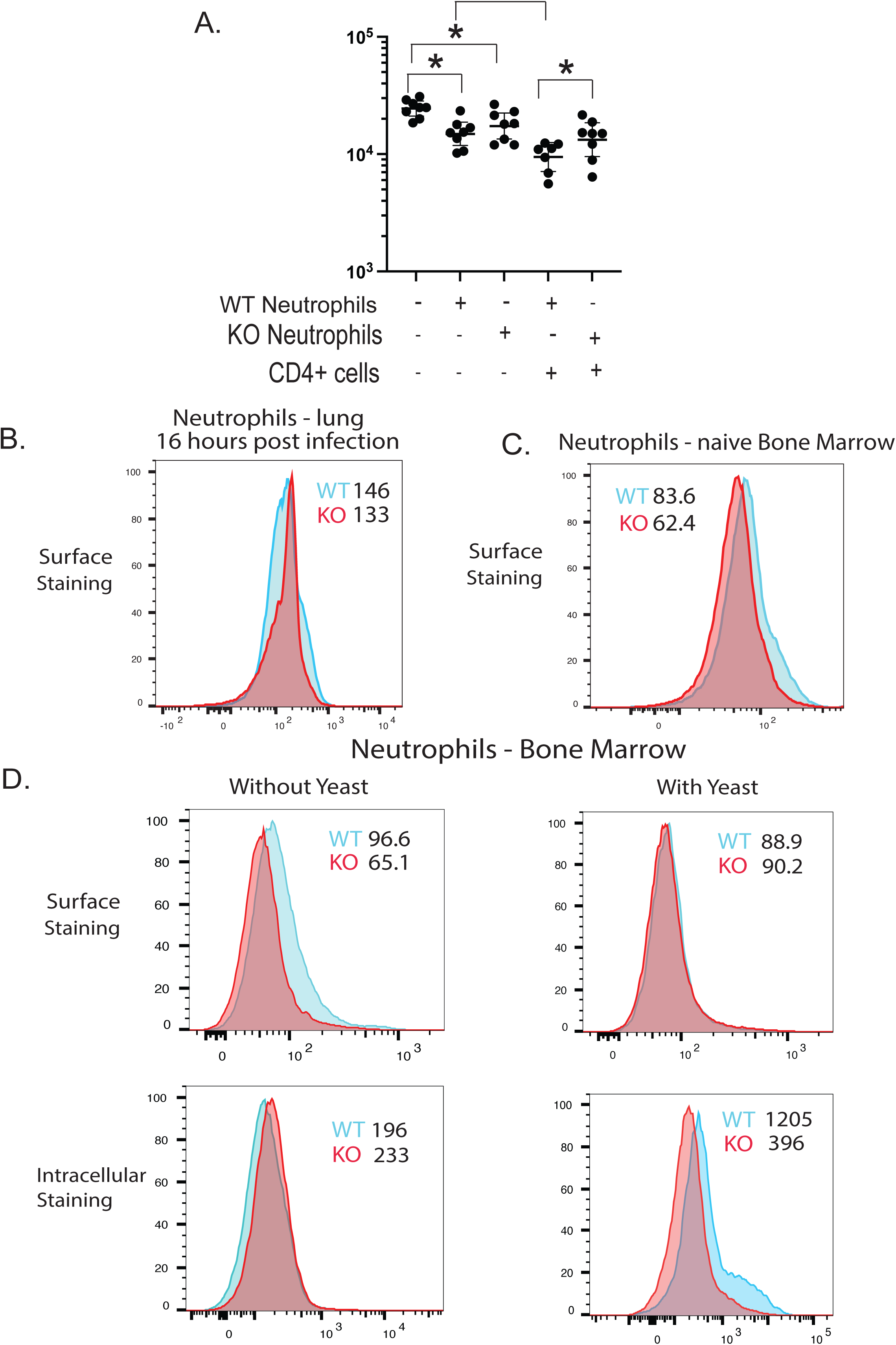
Augmented neutrophil killing requires SLAMF1 expression on neutrophils. *In vitro* killing assay with wild type or SLAMF1 KO neutrophils and CD4^+^ T cells from WT mice. Neutrophils from bone marrow of naive mice were cocultured overnight with *Bd* yeast and CD4^+^ T cells harvested from the lungs of WT mice 16 hrs post-infection **(A)**. 8 replicates plated for CFU. *p<0.05, two tailed Mann-Whitney T test. *Ex vivo* staining of neutrophils from lungs 16 hr post-infection **(B)** and from bone marrow of naive mice **(C)**. Surface and intracellular staining of neutrophils from the bone marrow with anti-SLAMF1 (CD150) after overnight coculture with or without *Bd* yeast **(D)** Geometric MFI of staining. Plots are concatenates from 5 mice/group.

In view of the above findings and conclusion, we were surprised that we saw no SLAMF1 expression on neutrophils from wild type mice compared to SLAMF1-deficient mice that were harvested from the lungs at 16 hours post-infection (**Fig. 5B**). However, our *in vitro* coculture results from above suggest SLAMF1 is required on neutrophils for leukocytes to augment killing. We hypothesized this difference in expression of SLAMF1 detected on the neutrophil may be attributed to contact with *Bd* yeast *in vivo* in the lung, as the *in vitro* assay uses naïve neutrophils from bone marrow. Thus, we observed naïve neutrophils from wild type mice harvested from the bone marrow expressed SLAMF1 (**Fig. 5C**). We hypothesized that SLAMF1 on neutrophils is down-regulated or internalized upon exposure to *Bd* yeast in the lung. To test this hypothesis, we set up a coculture with neutrophils and yeast and analyzed surface expression of SLAMF1 after yeast exposure or not. Neutrophils from wild type mice expressed SLAMF1 on the surface when cultured without yeast but not after overnight coculture with yeast (**Fig. 5D**). After coculture, neutrophils from wild type mice showed intracellular staining of SLAMF1 indicating that the receptor becomes internalized upon yeast exposure. These results are consistent with the idea that, early in infection, homotypic SLAMF1:SLAMF1 interactions between CD4^+^ T cells and neutrophils are required to augment neutrophil killing of yeast.

### Homotypic SLAMF1:SLAMF1 interactions between innate CD4^+^ T cells and neutrophils augment fungal killing by neutrophils via production of soluble factors

We investigated whether homotypic SLAMF1:SLAMF1 interactions between innate T cells and neutrophils may produce soluble factors that activate neutrophils to kill *Bd*. To test this idea, we established a transwell assay in which a lower well contained yeast, innate T cells and neutrophils (yielding homotypic SLAMF1:SLAMF1 interactions) and an upper well contained neutrophils and yeast. Soluble factors from the lower well could then diffuse across the membrane to activate neutrophils in the upper well (**Fig. 6A**). To verify that soluble factors are capable of passing through the 0.4 mm pores of the transmembrane, we used recombinant IFN-γ (17 kDA) known to activate neutrophils and increase their release of ROS and pro inflammatory cytokines [17] . We placed yeast and neutrophils in the top insert and added IFN-γ alone in the bottom insert. The addition of IFN-γ augmented neutrophil killing of the yeast in the top well (**Fig. 6B**) indicating that our transwell system allows soluble factors of at least 17 kDA to cross the membrane. The addition of CD4^+^ T cells in the bottom well augmented killing of yeast by neutrophils in the bottom well and in the corresponding top well (**Fig. 6C**). This result indicates that homotypic SLAMF1:SLAMF1 interactions between CD4^+^ T cells and neutrophils in the bottom well released soluble factors that activated neutrophils in the top well. From these data we propose a model by which augmented killing by neutrophils could be the consequence of SLAMF-1 mediated cell-contact or contact-dependent production of soluble factors or both cell contact and soluble factors (**Fig. 6D**).

**Figure 6.**
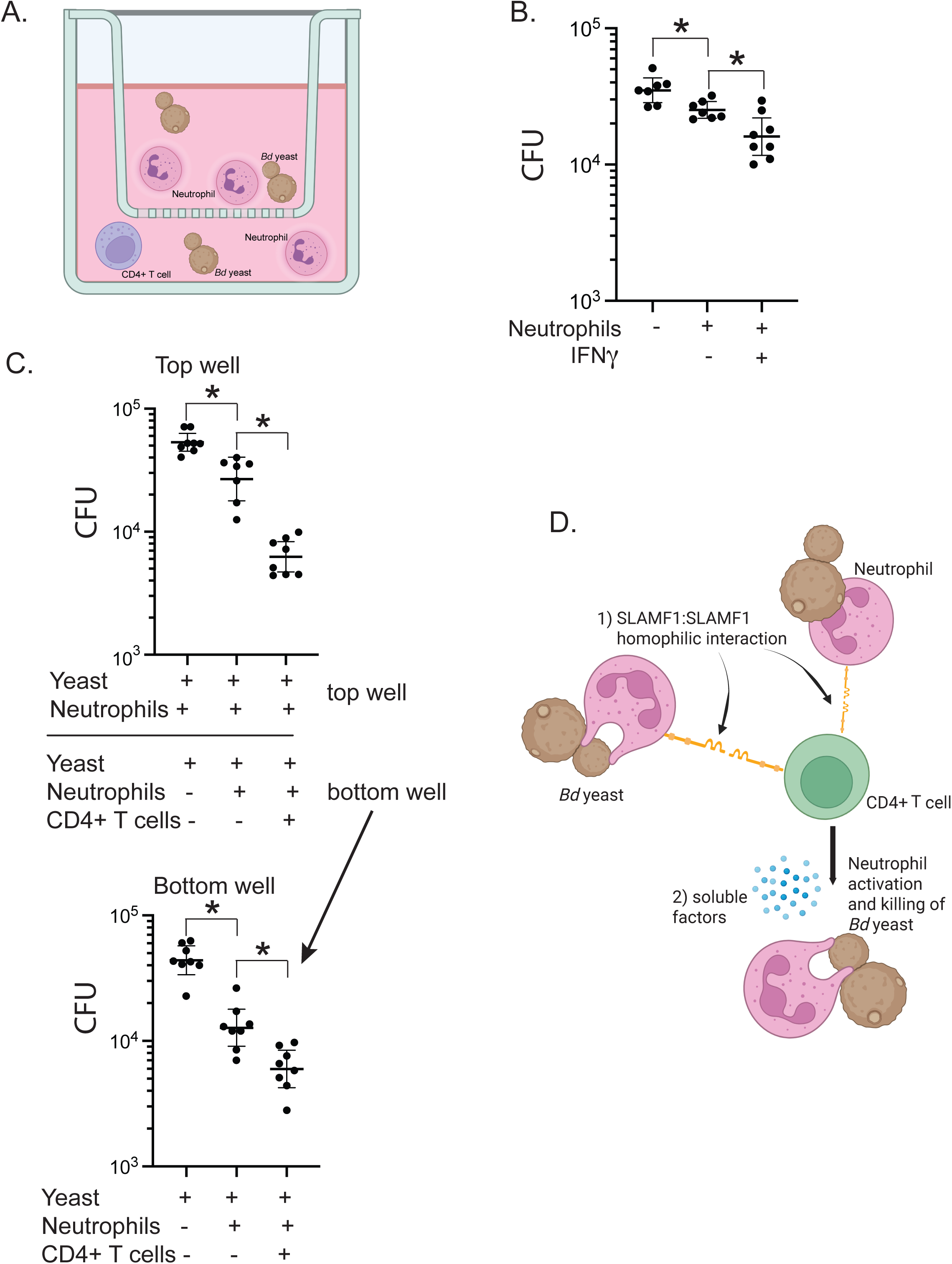
Transwell assay to investigate the role of soluble factors in SLAMF1-dependent neutrophil killing of *Bd*. Format of *in vitro* transwell assay: activating cells that potentially produce soluble factor in the bottom well; sensing neutrophils in the top well. *Bd* yeast and neutrophils from bone marrow of naive wild type mice were placed in the top insert wells. *Bd* yeast, neutrophils, and CD4^+^ T cells harvested from the lungs of wild type mice 16 hr post challenge were placed in the bottom well **(A)**. Validation of transwell system using IFN-γ as a positive control. Neutrophils and yeast were placed in the top well and recombinant IFN-γ added in the bottom well. 16 hours later, CFU were plated from the top well **(B).** Transwell system results after overnight coculture, 8 replicates plated for CFU (**C**). *p<0.05, two tailed Mann-Whitney T test. Model for SLAMF1 functions (**D**): 1) Cell contact through SLAMF1: SLAMF1 homotypic interactions between neutrophils and CD4^+^ T cells are requisite for neutrophil activation. 2) Homotypic SLAMF1:SLAMF1 interactions generate soluble factors that also augment neutrophil killing of *Bd* yeast.

## DISCUSSION

The role of SLAMF1 in mediating antifungal immunity is incompletely understood. Furthermore, little work has been done to address the mechanisms of SLAMF1 receptor interaction among innate immune cells using experimental models of microbial infection. Here, we investigate the role of SLAMF1 during the innate immune response to pulmonary fungal infection. We found that SLAMF1 is required to restrain pulmonary infection with *Bd*. Neutrophils and monocytes from SLAMF1 knockout mice killed yeast inefficiently *in vivo* in the lung in association with impaired ROS and NO production. Conversely, these cells killed *Bd* efficiently *in vitr*o pointing to a cell extrinsic role of SLAMF1 for phagocyte function *in vivo*. Our study therefore addressed the cellular and molecular mechanisms that extrinsically augment neutrophil killing in a SLAMF1 dependent manner.

We focused our study of extrinsic activators of neutrophil function by investigating lung innate CD4^+^ T cells, TCRγδ^+^ T cells and monocytes since they expressed SLAMF1 during pulmonary fungal infection. We generated conditional knockout mice that upon tamoxifen treatment lack SLAMF1 on these three candidate cells that might augment neutrophil function. *In vivo* and *in vitro* assays with cells from these mice enabled us to determine whether and how each cell subset participates in the SLAMF1 dependent activation of neutrophils.

*In vivo*, SLAMF1 expression by CD4^+^ T cells was indispensable to restrain fungal infection. *In vitro*, the addition of CD4^+^ T cells augmented neutrophil killing in a SLAMF1-dependent manner. Augmented killing by SLAMF1 expressing CD4^+^ T cells also required SLAMF1 expression on neutrophils, indicating that increased neutrophil activation was mediated by homotypic SLAMF1:SLAMF1 interaction between CD4^+^ T cells and neutrophils. Moreover, homotypic SLAMF1:SLAMF1 interactions between CD4^+^ T cells and neutrophils produced soluble factors that augmented neutrophil killing. Thus, augmented killing by neutrophils could have been the consequence of SLAMF-1 mediated cell-contact or contact-dependent production of soluble factors or both cell contact and soluble factors. This question is suitable for future investigation.

Protective immunity to the intracellular pathogen *Mycobacterium tuberculosis* (*Mtb*) also required cognate interactions between CD4^+^ T cells and macrophages[18]. However, contact dependency was not due to homotypic SLAMF1:SLAMF1 interactions but between MHC II on macrophages and TCR on antigen-specific CD4^+^ T cells which led to increased SLAMF1 expression that enhanced the generation of reactive oxygen species and restricted *Mtb* replication. In previous work we found that priming and function of antigen-specific CD4^+^ T cells that mediate protection against *B. dermatitidis* infection did not require SLAMF1[12].

We have previously shown that TCRγδ^+^ T cells and CD4^+^ T cells recruit and activate neutrophils and innate resistance to *Bd* infection [3]. In that study, the lack of TCRγδ^+^ T cells did not result in higher fungal burden but led to increased numbers of CD4^+^ T cells that likely compensated for the absence of TRCγδ^+^ T cells, since CD4^+^ T cells were found to be indispensable for host protection. We did not previously investigate the role of SLAMF1 in innate resistance mediated by CD4^+^ T cells or TCRγδ^+^ T cells. The mechanics that initially activate these innate populations at the site of infection have not been previously identified to our knowledge. It is possible that homotypic SLAMF1:SLAMF1 interactions between these innate T cells and neutrophils and associated release of soluble products are at the root of these responses during pulmonary fungal infection and possibly other mucosal infections with fungi, for example *Candida albicans*.

In a pulmonary model of aspergillosis, monocytes have been reported to exhibit two mechanisms by which they can exert innate antifungal activity [4]. First, CCR2^+^ monocytes and monocyte-derived DCs (Mo-DCs) conditioned the lung inflammatory milieu to augment neutrophil conidiacidal activity. Second, conidial uptake by monocytes temporally coincided with their differentiation into Mo-DCs, a process that resulted in direct conidial killing. In our study, using monocytes from conditional knockout mice that specifically lacked SLAMF1 only in monocytes demonstrated that the addition of monocytes to the coculture augmented killing in the wells in a SLAMF1 independent manner. These results suggest that monocytes also augment killing by neutrophils in a manner that does not require SLAMF1 to condition the neutrophils or that monocytes independently contributed to reduced CFU by directly killing the yeast.

In addition to neutrophils communicating with innate lymphocytes via homotypic SLAMF1:SLAMF1 interactions, other modes of SLAMF1 action remain possible, for example SLAMF1 on neutrophils might directly sense *Bd* yeast. This idea is supported by our observation that coculture of neutrophils and *Bd* yeast *in vitro* led to the internalization of SLAMF1. Consequently, we did not detect surface expression of SLAMF1 on lung neutrophils following infection with *Bd*. As part of SLAMF1 sensing, it is also possible that smaller *Bd* yeast (the size of the organism ranges from 8-20 mm) can be phagocytosed by neutrophils (range in size between 12-15mm) and, during this process, SLAMF1 is internalized. The potential sensing function of SLAMF1 is a topic of future investigation.

Instead of internalization, enhanced expression of SLAMF1 on human neutrophils has been reported after exposure to *Mtb* exposure. Stimulation with *Mtb* or its antigens *in vitro* induced surface SLAMF1 expression on neutrophils [19]. Diverse mycobacterial components were recognized by human neutrophils during stimulation with *Mtb*-antigens and induced SLAMF1 expression. In addition, increased SLAMF1 expression induced by *Mtb* involved the MAPKs MAPK14/p38 and MAPK/ERK and NADPH-dependent ROS production and autophagy. The authors of that study concluded that SLAMF1 is an innate receptor in human neutrophils that can regulate cellular responses against *Mtb* e.g. inducing autophagy in myeloid cells or increasing Th1 responses and thereby play an important role in human immunity against *Mtb*. Hence, SLAMF1 may be an attractive target for host directed therapy through increasing efficacy of pathogen control or reducing tissue damage due to infection.

Our study raises several important questions about SLAMF1 that have implications for innate immunity to fungi such as *Bd* and other intracellular microbes. For example, SLAMF1 as a sensor of *Bd* and the receptor’s potential role in phagocytosis; SLAMF1 induced signaling on innate lymphocytes and neutrophils; the identity of soluble factors released by homotypic SLAMF1:SLAMF1 interactions between neutrophils and innate T cells; and, lastly, the potential use of soluble SLAMF1 receptor for anti-fungal immunotherapy. Insights from such work will deepen our understanding of the diverse SLAMF1’s receptor functions that regulate innate immune interactions and provide the foundation for targeted host directed therapies.

## MATERIALS AND METHODS

### Fungi

*Blastomyces dermatitidis* strains used were wild-type, virulent *B*. *dermatitidis* ATCC strain 26199 and DsRed26199[20]. *B*. *dermatitidis* was grown as yeast on Middlebrook 7H10 agar with oleic acid-albumin complex (Sigma) at 39°C.

### Mice

SLAMF1 knockout mice were generated and obtained from Dr. Cox Terhorst at the Beth Israel Deaconess Medical Center at the Harvard Medical School in Boston[21]. C57BL/6 mice were obtained from Jackson Laboratory and bred at our facility. Heterozygous SLAMF1^fl/+^ were obtained from Shanghai Model Organisms and bred to homozygosity. Homozygous SLAMF1^fl/fl^ were crossed with cell specific cre mice (obtained from The Jackson Laboratory to generate conditional knockout mice. Cre mice were B6.FVB-Tg(EIIa-cre expressed in embryonic cells) C5379Lmgd/J (stock #003724), B6(129X1)-Tg(Cd4 cre/ERT2 expressed by CD4^+^ T cells)11Gnri/J (stock #22356), B6.129S-Tcrdtm1.1(cre/ERT2 expressed by TCRγδ^+^ T cells)Zhu/J (stock #031679) and C57BL/6-Ccr2em1(icre/ERT2 expressed by monocytes) Peng/J (stock #035229). Conditional CCR2-Cre^+^ x SLAMF1*^fl/fl^* mice were generated using *CCR2-CreER-GFP* mice (stock #035229) that express *cre-ER^T2^*and *EGFP* directed from the endogenous chemokine (C-C motif) receptor 2 (*Ccr2*) promoter.

### Generation of conditional SLAMF1 knockout mice

To generate conditional SLAMF1 knockout mice we first bred commercial SLAMF1^fl/+^ mice to homozygosity (**SFig. 3A**). To validate homozygous SLAMF1^fl/fl^ mice we crossed them with EIIa-cre mice[22], which carry a cre transgene under the control of the adenovirus EIIa promoter that targets expression of Cre recombinase to the early mouse embryo, and are useful for germ line deletion of *loxP*-flanked genes. If EIIa cre x SLAMF1^fl/fl^ mice ablate SLAMF1 at the germline level, they should lack SLAMF1 expression throughout the body and exhibit a similar resistance phenotype to *Bd* infection as seen with the original full body SLAMF1 knockout mice (**Fig. 1A+B**).

SLAM1 staining and FACS analysis showed that CD4^+^ T cells, TCRγδ^+^ T cells and monocytes from EIIa cre x SLAMF1^fl/fl^ mice lacked SLAMF1 expression similar to full body SLAMF1 knockout mice (**SFig. 3B**) and exhibited accelerated death and increased lung CFU upon infection with *Bd* (**SFig. 3C+D**). These results indicated that EIIa cre x SLAMF1^fl/fl^ mice phenocopied SLAMF1 knockout mice.

#### Lung CFU and survival

Mice were challenged with 2×10^4^ *B. dermatitidis* yeast. When mice became moribund lungs were harvested and plated for lung CFU or euthanized for generating a survival curve.

### Lung processing

Mice were challenged with 2×10^4^ 26199 yeast. Lungs were harvested in wash buffer and dissociated in Miltenyi MACs tubes, then digested with 5ml collagenase D solution (collagenase buffer (10mM HEPES, 5mM KCl, 2.1 mM MgCl_2_, 1.8 mM CaCl_2_, 241 mM NaCl/ml) with 1mg/ml Collagenase D, 1μg/mL DNAse) for 30 min at 37C. Digestion was stopped with 100μl of 0.5M EDTA (Invitrogen 15575-038). Digested lungs were resuspended in 5 ml of 40% Percoll in wash buffer; 3 ml of 66% Percoll in PBS was underlaid (17-0891-01; GE Healthcare). Samples were spun for 20 min at 550 RCF at 22C. Lymphocytes were then harvested from the buffy coat layer and resuspended in desired media.

### Isolation of target cells from the lung and bone marrow

CD4^+^ and TCRγδ^+^ T cells were isolated from the lungs of infected mice (16 hours post-challenge) following lung processing protocol referenced above to harvest leukocytes, then according to the manufacture’s isolation protocols (Miltenyi Isolation kits (CD4+T cells; Cat#130-104-454 and TCRγδ; Cat# 130-092-125). Monocytes were isolated from the bone marrow following the manufacturer’s protocol (Monocyte: Cat# 130-100-629). Cell populations before and after isolation and negative fractions were stained for SLAMF1 and the purity assayed by FACS.

### In vitro killing assay

Neutrophils were used as effector cells for killing the yeast and harvested from bone marrow of wild type C57BL/6 mice. Bone marrow was flushed with PBS and aspirated to disaggregate large bone marrow pieces from hind legs and filtered through a 70 mM cell strainer (Fisher Cat# 22362548). Suspension was pelleted then resuspended in 6 mL of 0.2% NaCl to lyse red blood cells (RBCs). After 10 seconds, osmolarity was restored with the addition of 14 ml of 1.2% NaCl. Cell suspension was strained with a 70 mM cell strainer and pelleted. Pellet was resuspended in 5 ml of wash buffer and pipetted on top in a tube of 5mL 62% Percoll in PBS with 2 ml of 83% Percoll in wash buffer underlaid. Samples were spun for 30 min at 1000 RCF at 22C. Neutrophils were then harvested from buffy coat layer between 62% and 83% Percoll, washed with wash buffer, and resuspended to 4.5×10^6^/ml in complete RPMI (cRPMI, RPMI with 10% FBS, 1% penicillin and streptomycin). Yeast were resuspended in cRPMI to a concentration of 0.5×10^6^/ml. 100 μl of yeast, neutrophils, and target cell types were seeded in an untreated 96 well plate (Avantor #10861-561). Isolated cells were concentrated to a range of 5×10^6^ to 1×10^7^/mL for CD4^+^ T, 1×10^6^ to 1.5×10^6^/mL for TCRγδ^+^ T cells, and 1×10^6^ to 2×10^6^/mL for monocytes. For yeast only or yeast and neutrophil groups, wells were filled to a final volume of 300 μl with cRPMI. Samples were incubated for 16 hours before plating for CFU.

### In vitro transwell assay

Neutrophils were isolated as described in the *in vitro* killing assay. Neutrophils were concentrated to 4.5×10^6^/ml and 26199 yeast concentrated to 0.5×10^6^/ml. 50μl of yeast, neutrophils, and target cell types were placed in the upper well of a 96 transwell system with a 0.4mm polycarbonate membrane (Corning 3381) to allow for homotypic interactions. For yeast only or yeast and neutrophil groups, the upper well was filled to a volume of 150 μl with cRPMI. For positive control group, human recombinant IFN-g (R&D Systems 285-IF-100) in cRPMI was added to the top well for a final concentration of 150U/ml.[23] The bottom well was seeded with 100 μl of yeast and effector neutrophils. For the yeast only control group, cRPMI was added for a final volume of 200 μl. The transwell assay was incubated at 37C for 16 hours. Yeast from the bottom well were plated to determine CFU.

### In vivo killing assay

To assess the ability of phagocytes to kill yeast *in vivo*, we employed Ds-Red 26199 yeast that loses red fluorescence in dead yeast as described [15,20]. Briefly, yeast were concentrated to 5×10^6^/ml and 0.5 ml was stained with 10μl of 1mg/ml Uvitex in the dark at 22C. After washing the yeast, the mice were challenged with 10^5^ Uvitex stained Ds-Red yeast. FMO controls were included by challenging mice with unstained 26199 yeast, Ds-Red yeast, and Uvitex stained 26199 yeast. Lungs were harvested 16 hours after challenge in wash buffer (RPMI with 1% FBS, 1% penicillin and streptomycin) and dissociated in Miltenyi MACs tubes and digested with 5ml collagenase D solution (collagenase buffer with 1mg/ml Collagenase D, 1ug/ml DNAse) for 25 min at 37C. Digestion was stopped with 100 μl of 0.5M EDTA (Invitrogen 15575-038) and RBCs were lysed for 4 min with 5ml ACK lysis buffer (Gibco A10492-01) then diluted with 20ml of wash buffer. Cells were washed and resuspended to a concentration of 2×10^7^/ml for staining. Samples were stained with LIVE/DEAD Fixable Near-IR Dead Cell Stain Kit (L34975; Invitrogen) and Fc block for 10 min at room temperature. Samples were surface stained at 4C for 30 minutes, following by fixation with 2% PFA (16% PFA diluted with ddH_2_O; VWR Cat# 28908). Panel included a dump channel to reduce non-specific staining. The antibody cocktail consisted of Siglec F BB515 (E50-2440; Cat # 564514; BD), Ly6G PerCP-Cy5.5 (1A8; Cat #127616; Biolegend), CD11c PE-Cy7 (N418; Cat# 117317; Biolegend), Ly6C BV785 (HK1.4; Cat# 128041; Biolegend), MHCII A647 (M5/114.15.2; Cat# 107618; Biolegend), TCRβ BUV395 (H57-597; Cat# 569248; BD), B220 BUV395 (RA3-6B2; Cat#563793; BD), Nk1.1 BUV395 (PK136; Cat# 564144; BD), CD11b BV737 (M1/70; Cat# 612800; BD).

### ROS and NO staining

Sample aliquots from *in vivo* killing assay were taken at staining step to be stained for ROS and NO. Samples were placed in a 96 well plate. For ROS staining the samples were stained with 10mg/ml DHR-123 (Invitrogen D23806) and incubated for 3 hours at 37C then surface stained following *in vivo* killing assay protocol. For the NO staining the samples were stained with 10 mM DAF-FM diacetate (Invitrogen D23844) and incubated for 10 to 20 minutes at 37C then washed with PBS and incubated for another 15 minutes. Cells were surfaced stained following *in vivo* killing assay protocol.

### Ex vivo SLAMF1 staining - Lungs

Leukocytes from the percoll gradient were stained with LIVE/DEAD Fixable Near-IR Dead Cell Stain Kit (L34975; Invitrogen) and Fc block for 10 min at room temperature. Samples were surface stained at 4C for 30 minutes, following by fixation with 2% PFA (16% PFA diluted with ddH_2_O; VWR Cat# 28908). The antibody staining panel consisted of SLAMF1 FITC (mShad150; Cat# 11-1502-82; Invitrogen) or BV785 (TC15-12F12.2; Cat# 115937; Biolegend), TCRγδ PerCP-Cy5.5 (GL3; Cat# 118118; Biolegend), Siglec F PerCP-Cy5.5 (E50-2440; Cat# 565526; BD), IL33Ra PE (DIH9; Cat# 145304; Biolegend), CD64 PE (X54-5/7.1; Cat# 139304; Biolegend), CD19 PE Dazzle (6D5; Cat# 115554; Biolegend), TCRβ PE-Cy7 (H57-597; Cat# 109222; Biolegend), CD11c PE-Cy7 (N418; Cat# 117317; Biolegend), NK-1.1 BV421 (PK136; Cat# 108741; Biolegend), CD103 eFluor450 (2E7; Cat# 48-1031-82; Invitrogen), Ly6C BV510 (HK1.4; Cat# 128033; Biolegend), CD8a BV650 (53-6.7; Cat# 100742; Biolegend), CD11b BV650 (M1/70; Cat# 101239; Biolegend), CD90.2 BV785 (30-H12; Cat# 105331; Biolegend) or BUV737 (53-2.1; Cat# 741701; BD), CD196 AF647 (140706; Cat# 557976; BD), MHCII AF700 (M5/114.15.2; Cat# 107622; Biolegend), CD4 BUV395 (GK1.5; Cat# 563790; BD), Ly6G BUV395 (1A8; Cat# 563978; BD).

### Surface and intracellular staining of SLAMF1 of neutrophils from the bone marrow

Bone marrow was harvested as described above and stained with the LIVE/DEAD Fixable Near-IR Dead Cell Stain Kit and Fc block for 10 min at room temperature. Samples were surface stained at 4C for 30 minutes using the SLAMF1 antibody panel as described above, washed with FACS buffer, then incubated in 100 μl of Cytofix/Cytoperm (BD Cat# 554714) at 4C for 20 minutes. Samples were washed twice with Perm/Wash buffer (BD Cat# 554714) and stained intracellularly with anti-CD150 BV-785 (TC15-12F12.2; Cat# 115937; Biolegend), washed again with Perm/Wash buffer, then fixated with 2% PFA (16% PFA diluted with ddH_2_O; VWR Cat# 28908). We also stained neutrophils that were exposed to yeast overnight. 4.5×10^5^ neutrophils and 5×10^4^ yeast in cRMPI were incubated for 16 hours in a 96 well plate prior to staining.

### Flow Cytometry

Samples were acquired on an LSR Fortessa and Cytek Aurora Spectral Cytometer, at the University of Wisconsin Carbone Cancer Center Flow Lab.

### Ethics Statement

Animal studies adhered to protocol M005891 approved by the IACUC of UW-Madison. Animal studies were compliant with provisions established by the Animal Welfare Act and the Public Health Services (PHS) Policy on the Humane Care and Use of Laboratory Animals.

### Statistics

All statistics were calculated in Prism 10 for Mac OS X, version 6.1. Differences in fungal burden (expressed as CFU) between two groups were analyzed by the Mann-Whitney U test for ranking data. In some instances, 1-way ANOVA was used when comparing multiple groups, and when a result was significant, a Tukey’s or Dunnett’s post hoc test was used to adjust for multiple comparisons. For comparison of fungal burden among three or more groups of mice, the Kruskal-Wallis test, a nonparametric ranking method, was used. Survival data were examined by the Kaplan-Meier test using log rank analysis to compare survival plots as reported previously (6). A p value of ≤0.05 was considered statistically significant. Comparisons in many experiments yielded p values at or below the value of p = 0.05, however we consistently used only one asterisk throughout to denote any statistically significant difference regardless of the exact value below 0.05.

**Supplementary Figure 1 to main Figure 1.**
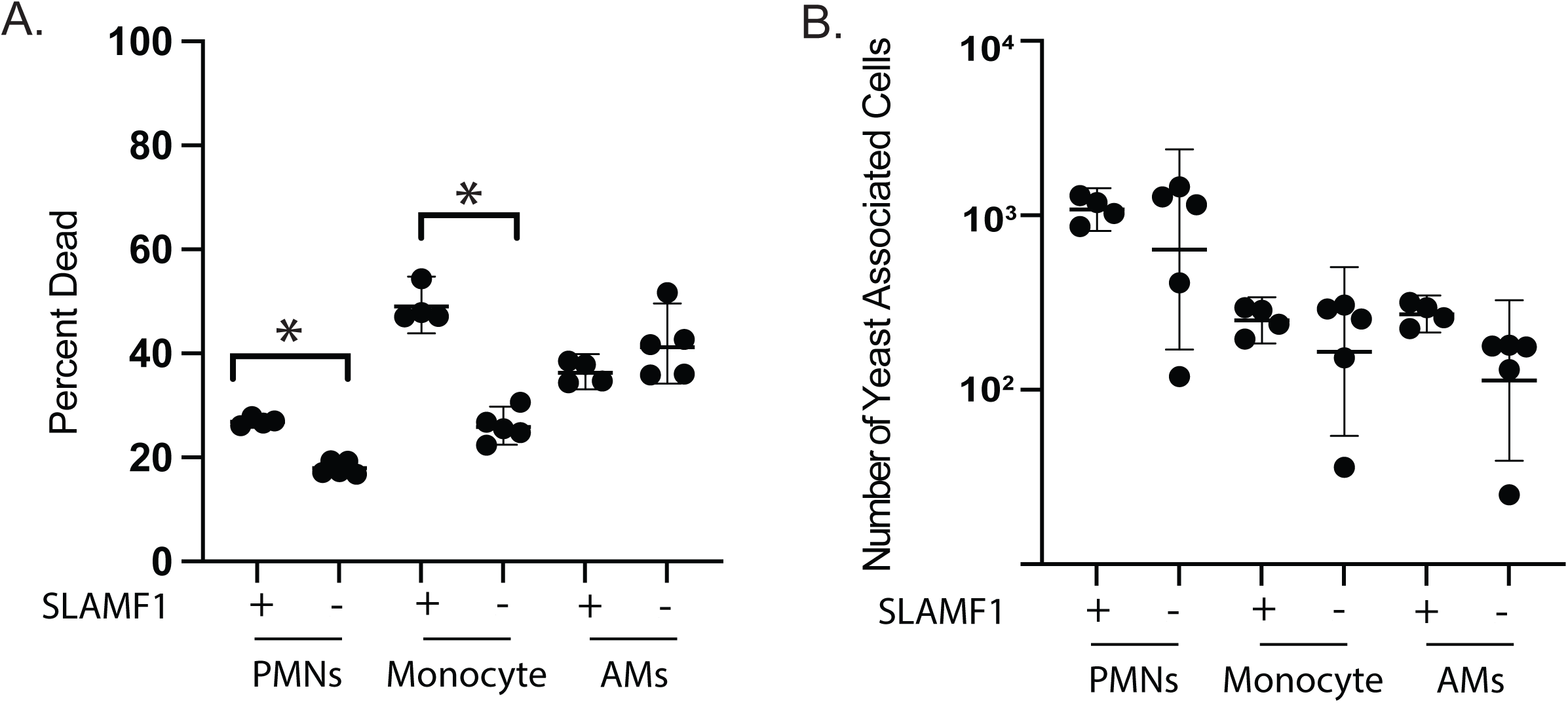
*In vivo* killing assay: Percent of lung phagocytes associated with dead yeast (**A**) and number of alveolar macrophages, neutrophils, and monocytes associated with total number of yeast for wild type and SLAMF1 knockout mice (**B**). *p<0.05, two tailed Mann-Whitney T test.

**Supplementary Figure 2 to main Figure 3.**
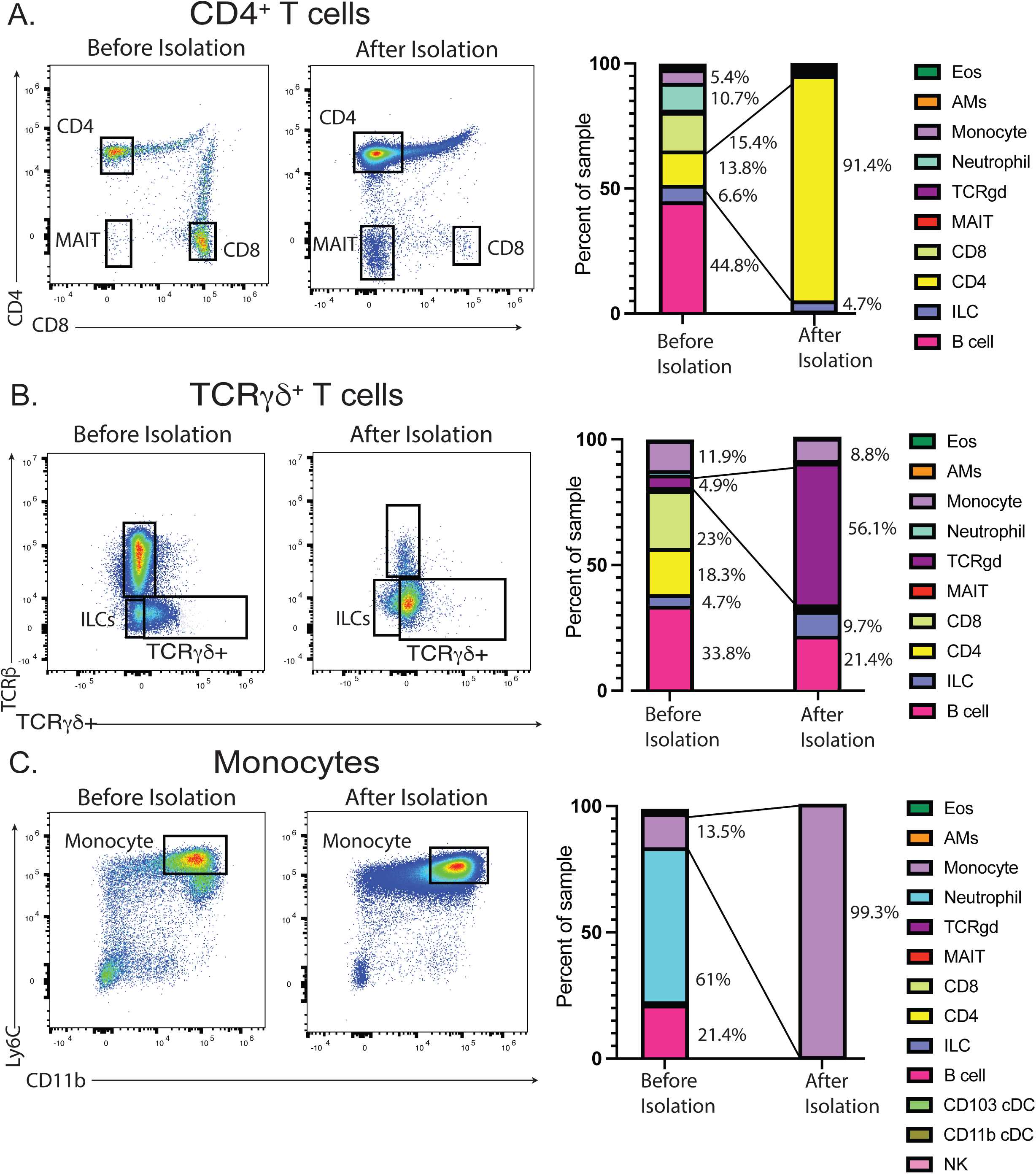
Flow analysis and stacked bar graphs of lung leukocyte populations before and after isolation of CD4^+^ T cells **(A)** and TCRγδ^+^ T cells **(B)**, and analysis of bone marrow before and after isolation of monocytes **(C).** To identify cell types of interest before and after enrichment we used a panel of multiplexing antibodies to identify 17 pulmonary leukocyte subsets with 12 color flow cytometry [16]. Stacked bar graphs show percentage of each immune cell type relative to all immune cells identified by the Lymphoid Myeloid panel.

**Supplementary Figure 3 to main Figure 4.**
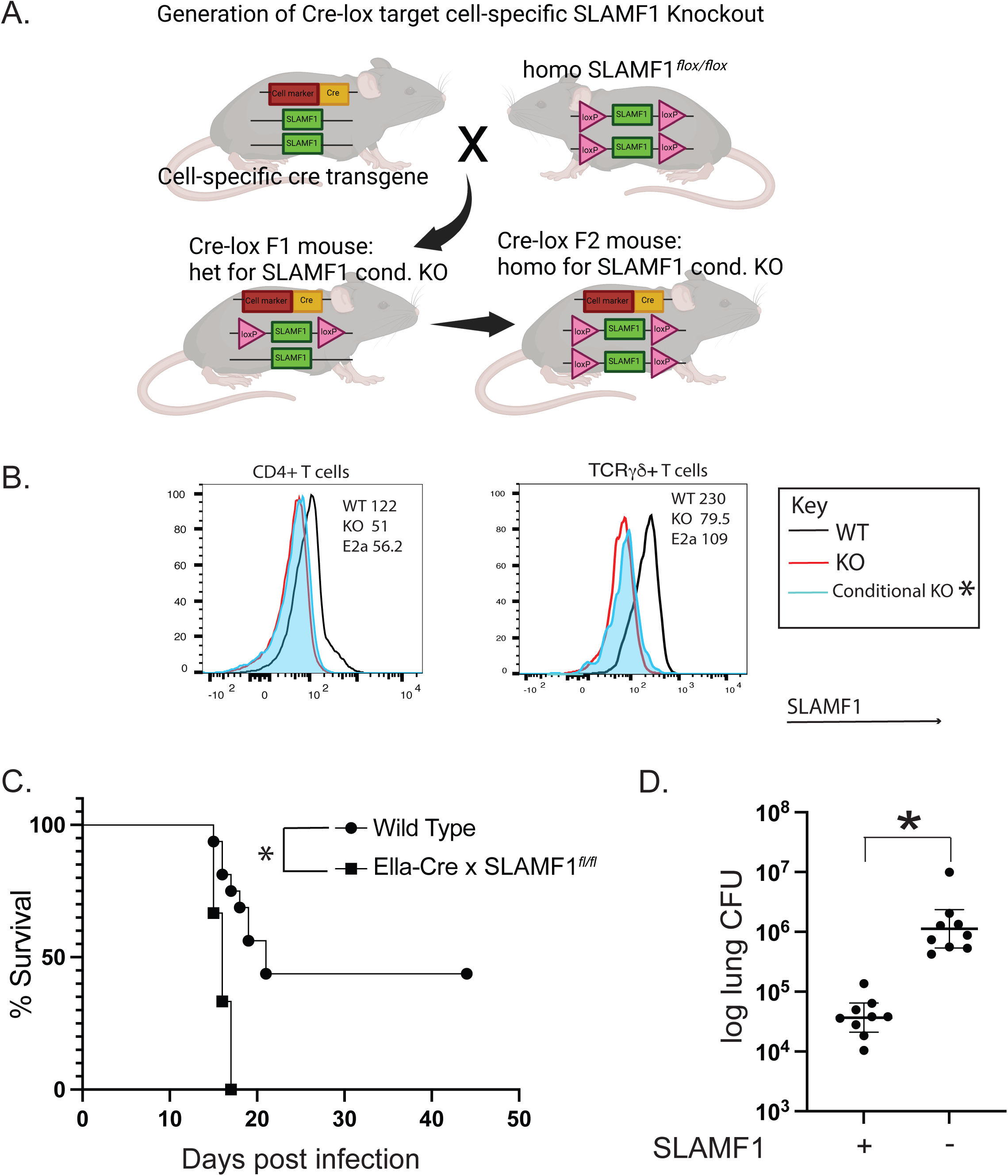
Breeding scheme for the generation of conditional SLAMF1knockout mice. To validate commercial SLAMF1 floxed mice we bred them first to homozygosity and crossed them with Ella-Cre mice to generate Ella-cre x SLAMF1^fl/fl^ mice that lack SLAMF1 in embryonic cells **(A).** SLAMF1 expression on target cells for Ella-cre x SLAMF1^fl/fl^ mice in comparison to wildtype and SLAMF1 KO mice. Geometric MFI of staining. Plots are concatenates from 5 mice/group. *p<0.05 vs WT, two tailed Mann-Whitney T test **(B).** Survival **(C)** and lung CFU **(D)** 9 days post infection of Ella-cre x SLAMF1^fl/fl^ mice and WT mice infected with *Bd*. *p<0.05, Kaplan Meier test for survival. CFU from at least 9 mice/group are expressed as Log_10_ plotted with geometric mean ± geometric SD *p<0.05 two tailed Mann-Whitney T test.

**Supplementary Figure 4 to Figure 4.**
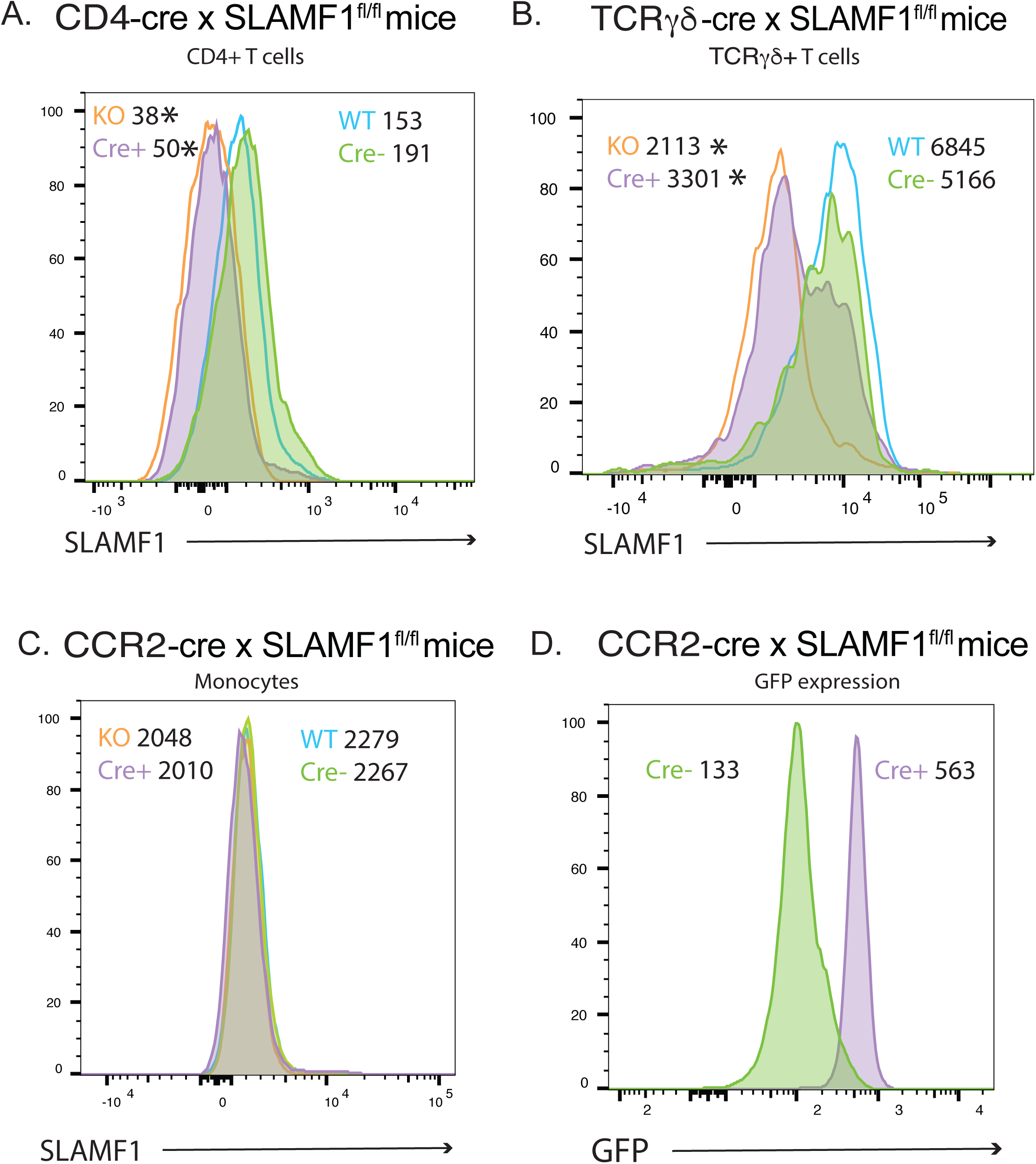
*Ex vivo* staining of lung leukocytes for SLAMF1 in CD4-cre x SLAMF1^fl/fl^ mice **(A)** and TCRγδ-cre x SLAMF1fl/fl mice **(B)**. *Ex vivo* staining of bone marrow for SLAMF1 in CCR2-cre x SLAMF1*^fl/fl^* mice **(C).** Plots show the histograms and geometric mean fluorescent intensity (MFI) of SLAMF1 staining for corresponding target cells in Cre^+^ and Cre^-^ mice in comparison to wild type and SLAMF1 knockout mice. GFP expression by conditional CCR2-Cre^+^ x SLAMF1*^fl/fl^* mice that were generated using *CCR2-CreER-GFP* mice (**D**). Plots are concatenates from 5 or more mice/group. *p<0.05 vs SLAMF1 expressing cells (WT vs KO, Cre^-^ vs Cre^+^), two tailed Mann-Whitney T test.

## Acknowledgement

The work was supported by NIH grants R21AI173718 (MW), R01AI040996 (BK/MW), R37 AI035681 (BK), and R01 AI168370 (BK). Flow samples were processed at the University of Wisconsin Carbone Cancer Center (UWCCC) Flow Core Facility on a BD LSR Fortessa and Aurora that was purchased with the NIH shared instrumentation grant 1S100OD018202-01 and University of Wisconsin Carbone Cancer Center Support grant P30 CA014520.

## REFERENCES

1. Wüthrich M, Deepe GS, Jr., Klein B (2012) Adaptive immunity to fungi. Annu Rev Immun30: 115–148.

2. Roy RM, Klein BS (2012) Dendritic cells in antifungal immunity and vaccine design. Cell Host Microbe 11: 436–446.

3. Hernandez-Santos N, Wiesner DL, Fites JS, McDermott AJ, Warner T, et al. (2018) Lung Epithelial Cells Coordinate Innate Lymphocytes and Immunity against Pulmonary Fungal Infection. Cell Host Microbe 23: 511–522 e515.

4. Espinosa V, Jhingran A, Dutta O, Kasahara S, Donnelly R, et al. (2014) Inflammatory monocytes orchestrate innate antifungal immunity in the lung. PLoS Pathog 10: e1003940.

5. Caffrey AK, Lehmann MM, Zickovich JM, Espinosa V, Shepardson KM, et al. (2015) IL-1alpha signaling is critical for leukocyte recruitment after pulmonary Aspergillus fumigatus challenge. PLoS Pathog 11: e1004625.

6. Romero X, Sintes J, Engel P (2014) Role of SLAM family receptors and specific adapter SAP in innate-like lymphocytes. Crit Rev Immunol 34: 263–299.

7. Chen S, Li D, Wang Y, Li Q, Dong Z (2020) Regulation of MHC class I-independent NK cell education by SLAM family receptors. Adv Immunol 145: 159–185.

8. Veillette A (2006) Immune regulation by SLAM family receptors and SAP-related adaptors. Nat Rev Immunol 6: 56–66.

9. Tatsuo H, Ono N, Tanaka K, Yanagi Y (2000) SLAM (CDw150) is a cellular receptor for measles virus. Nature 406: 893–897.

10. van Driel BJ, Liao G, Engel P, Terhorst C (2016) Responses to Microbial Challenges by SLAMF Receptors. Front Immunol 7: 4.

11. Kohn EM, dos Santos Dias L, Dobson HE, He X, Wang H, et al. (2022) SLAMF1 is dispensable for vaccine induced T cell development but required for resistance to fungal infection. J Immunol In press.

12. Kohn EM, Dos Santos Dias L, Dobson HE, He X, Wang H, et al. (2022) SLAMF1 Is Dispensable for Vaccine-Induced T Cell Development but Required for Resistance to Fungal Infection. J Immunol 208: 1417–1423.

13. Sterkel AK, Mettelman R, Wuthrich M, Klein BS (2015) The unappreciated intracellular lifestyle of Blastomyces dermatitidis. J Immunol 194: 1796–1805.

14. Fites JS, Gui M, Kernien JF, Negoro P, Dagher Z, et al. (2018) An unappreciated role for neutrophil-DC hybrids in immunity to invasive fungal infections. PLoS Pathog 14: e1007073.

15. Wang H, Lee TJ, Fites SJ, Merkhofer R, Zarnowski R, et al. (2017) Ligation of Dectin-2 with a novel microbial ligand promotes adjuvant activity for vaccination. PLoS Pathog 13: e1006568.

16. Wiesner DL, Merkhofer RM, Ober C, Kujoth GC, Niu M, et al. (2020) Club Cell TRPV4 Serves as a Damage Sensor Driving Lung Allergic Inflammation. Cell Host Microbe 27: 614–628 e616.

17. Marchi LF, Sesti-Costa R, Ignacchiti MD, Chedraoui-Silva S, Mantovani B (2014) In vitro activation of mouse neutrophils by recombinant human interferon-gamma: increased phagocytosis and release of reactive oxygen species and pro-inflammatory cytokines. Int Immunopharmacol 18: 228–235.

18. Krishna Prasad GVR, Grigsby SJ, Erkenswick GA, Portal-Celhay C, Mittal E, et al. (2025) Macrophage-T cell interactions promote SLAMF1 expression for enhanced TB defense. Nat Commun 16: 6794.

19. Pellegrini JM, Sabbione F, Morelli MP, Tateosian NL, Castello FA, et al. (2021) Neutrophil autophagy during human active tuberculosis is modulated by SLAMF1. Autophagy 17: 2629–2638.

20. Sterkel AK, Lorenzini JL, Fites JS, Subramanian Vignesh K, Sullivan TD, et al. (2016) Fungal Mimicry of a Mammalian Aminopeptidase Disables Innate Immunity and Promotes Pathogenicity. Cell Host Microbe 19: 361–374.

21. Wang H, LeBert V, Hung CY, Galles K, Saijo S, et al. (2014) C-type lectin receptors differentially induce th17 cells and vaccine immunity to the endemic mycosis of North America. The Journal of Immunology 192: 1107–1119.

22. Lakso M, Pichel JG, Gorman JR, Sauer B, Okamoto Y, et al. (1996) Efficient in vivo manipulation of mouse genomic sequences at the zygote stage. Proc Natl Acad Sci U S A 93: 5860–5865.

23. Marchi LF, Sesti-Costa R, Ignacchiti MDC, Chedraoui-Silva S, Mantovani B (2014) In vitro activation of mouse neutrophils by recombinant human interferon-gamma: increased phagocytosis and release of reactive oxygen species and pro-inflammatory cytokines. International immunopharmacology 18: 228–235.

